# Dynamics and Activation of Membrane-Bound B Cell Receptor Assembly

**DOI:** 10.1101/2024.07.10.602784

**Authors:** Hung N. Do, Mingfei Zhao, S. Munir Alam, S. Gnanakaran

## Abstract

B-cell receptor complexes (BCR) are expressed on the surface of a B-cell and are critical in antigen recognition and modulating the adaptive immune response. Even though the relevance of antibodies has been known for almost a hundred years, the antigen-dependent activation mechanism of B-cells has remained elusive. Several models have been proposed for BCR activation, including cross-linking, conformation-induced oligomerization, and dissociation activation models. Recently, the first cryo-EM structures of the human B-cell antigen receptor of the IgM and IgG isotypes were published that validated the asymmetric organization of the BCR complex. Here, we have carried out extensive molecular dynamics simulations to probe the conformational changes upon antigen binding and the influence of the membrane lipids. We identified two critical dynamical events that could be associated with antigen-dependent activation of BCR. First, antigen binding caused increased flexibility in regions distal to the antigen binding site. Second, antigen binding altered the rearrangement of IgM transmembrane helices, including the relative interaction of Igα/Igβ that mediates intracellular signaling. Furthermore, these transmembrane rearrangements led to changes in localized lipid composition.

## Introduction

B-cell receptors (BCRs) stand sentry on the front lines of the body’s defenses against infection, functioning by binding foreign substances known as antigens and activating the adaptive immune system^1^. On naïve B cells, one BCR class is a complex of *IgM* containing dimer of antigen-binding the Fab domains, Fc domains, transmembrane helices (TM), and the Igα/Igβ heterodimer signaling component^1^. Despite BCRs’ importance in adaptive immunity, mechanistic details of antigen-dependent activation remain elusive. However, the BCRs are known to be activated and clustered on the plasma membrane upon antigen binding^2,3^, which also triggers signal transduction by Igα/Igβ^3,4^. A recent study also showed that BCRs’ activation depends on the lipid membrane domains, and activated BCRs prefer more ordered domains in the plasma membrane vesicles^5^.

Several different models have been proposed for the BCR activation mechanism^2,4,6,7^. The first model, the cross-linking model, assumed the initiation of BCR signaling resulted from the aggregation of monomeric BCRs as multivalent antigens were observed to dominantly activate B cells^7–10^. This model drew attention because it proposed no signal needed to be propagated from the extracellular domain to the cytoplasm of the BCR^7–10^. Instead, the aggregation of the Igα-Igβ intracellular domains caused the cross-phosphorylation of the ITAMs motifs (the immunoreceptor tyrosine-based activation motif) by associated tyrosine kinases^7–10^. Two alternative models, namely the conformation-induced oligomerization model and dissociation activation model, have been proposed since the classical cross-linking model was considered too simplistic to account for diverse types of antigens that bound to the BCRs^4,7,11–15^. Based on the conformation-induced oligomerization model, the binding of membrane antigens activated the ectodomains of the BCRs and exposed an oligomerization interface in the membrane proximal region (MPR)^11,12^. With the exposure of this interface, the BCRs oligomerized, leading to perturbations of the local lipid environment, opening of the cytoplasmic domains, and initiation of signaling^11,12^. In the dissociation activation model, antigen binding dissociated the auto-inhibitory clusters of the resting BCRs on the B-cell surface, causing the opening of the Igα/Igβ heterodimer intracellular ends to expose their ITAM phosphorylation motifs^7,13–15^.

Recent cryo-EM shows that BCR is an asymmetric complex where the membrane-bound immunoglobulin molecule (IgM) binds to a single Igα/Igβ heterodimer with a 1:1 stoichiometry^3^. This contrasts with the previous conventional (textbook) belief that it is a symmetric molecule where it forms an assembly with two Igα/Igβ heterodimers in a 1:2 complex^16^ and supports the biochemical/biophysical evidence of the BCR asymmetry^17,18^. In the BCR complex, the antigen-sensing homo-dimeric membrane-bound immunoglobulin is associated with signal-transducing heterodimeric Igα/Igβ in a 1:1 stoichiometry^3,19^. Unlike previous assumptions of a symmetric model, asymmetry arises due to the association of the signaling domain with a single TM domain (TMD) of IgM, leaving the other TMD vacant^3,19^. Also, the orientation of TMD prevents the association with another Igα/Igβ herterodimer^3,19^. Notably, one Ig domain is locked between the Igα and Igβ in the juxtamembrane region^3^. This organization makes the BCR slightly titled in the membrane and eliminates the possibility of a symmetric BCR complex as the ectodomains of the second Igα-Igβ signaling component would clash with the membrane lipid head groups^3^. Given this confirmation of the structural arrangement of the BCR complex, we have carried out extensive molecular dynamics (MD) simulations to evaluate the dynamics of this asymmetric complex. We consider both all-atom (AA) and coarse-grained (CG) MD simulations of a human B-cell antigen receptor of the IgM isotype with the specificity of the anti-HIV-1 CD4 binding site CH31 antibody^20,21^ to profile the different protein conformations of this assembly in the presence/absence of the antigen, HIV-1 envelope (Env) protein (including the gp120 envelope protein), in a complex membrane, which included 63.2% phosphatidylcholine (POPC), 12.6% phosphatidylethanolamine (POPE), 17.4% palmitoyl sphingomyelin (PSM), 0.5% ceramide (CER3), 2.2% diacylglycerol (DAGL), and 4.2% cholesterol (CHOL), based on the experimental percentage of displayed lipid types of the plasma membrane determined from a previous study^22^ (**Figure 1**). We uncover never-before-seen molecular allosteric alterations in dynamics when the antigen is bound to the BCR complex. Our MD based BCR model will contribute to understanding the antigen-dependent activation mechanism of BCRs.

**Figure 1.**
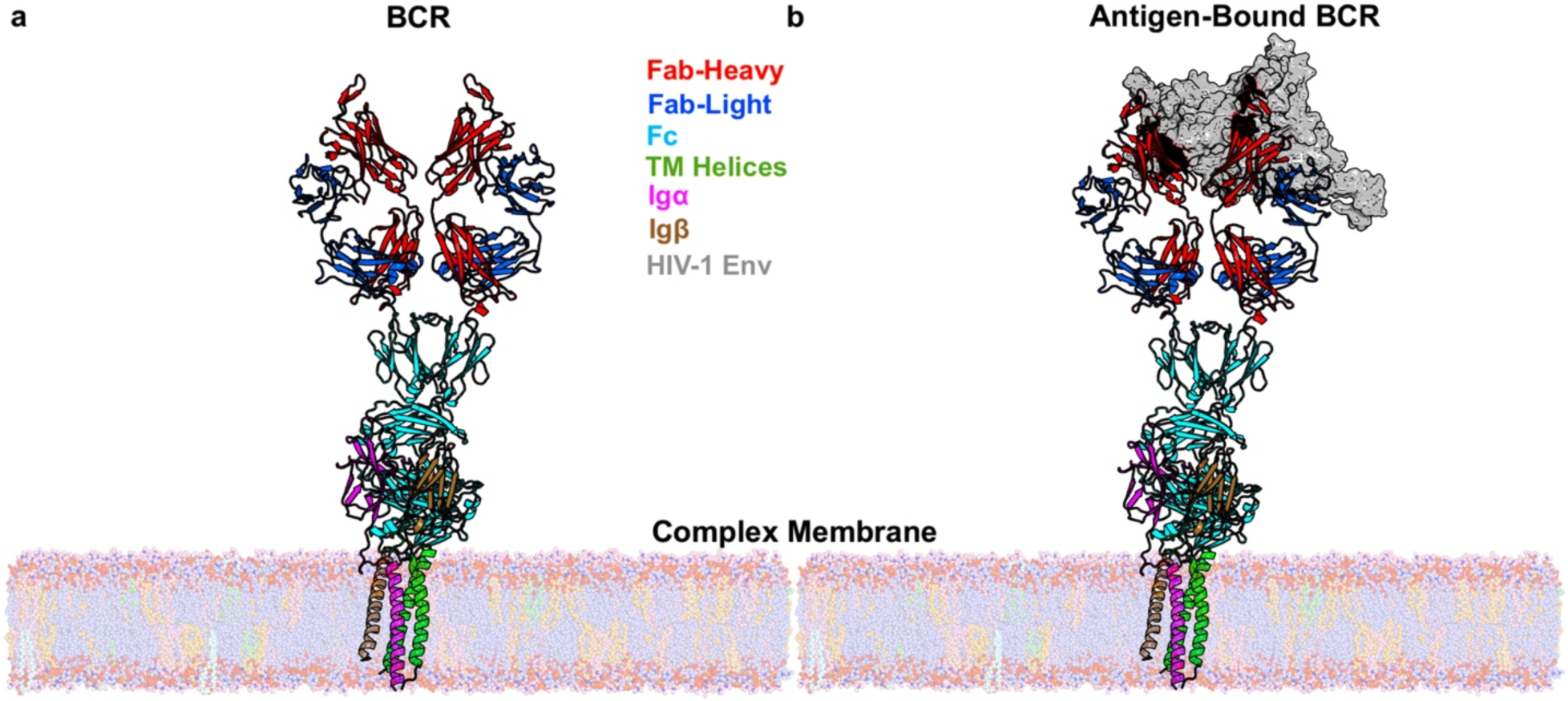
Simulation systems of the model B cell receptor (BCR) complexes. **(a)** The CH31 BCR in the complex membrane. **(b)** The CH31 BCR bound by a monomer of HIV-1 envelope protein (HIV-1 Env) in the complex membrane. The Fab heavy chains are colored red, Fab light chains are colored blue, Fc domains are colored cyan, transmembrane helices of BCR are colored green, Igα is colored magenta, Igβ is colored brown, and the HIV-1 Env is colored gray. The POPC molecules are colored light blue, POPE are colored light orange, PSM are colored light pink, diacylglycerols (DAGL) are colored pale green, cholesterols (CHOL) are colored wheat, and ceramides (CER3) are colored pale cyan.

## Results

### Description of BCR complex and antigen

The initial model for the BCR complex was based on the cryo-EM structure of the human B-cell antigen receptor of the *IgM* isotype with the CD4 binding-site antibody VRC01 Fab domain (PDB: 7XQ8)^3^. The BCR structure is a “Y”-shaped complex of *IgM* containing dimer of the VRC01 Fab domains connected to the Fc domains, and the TM helices, which associate to the Igα/Igβ heterodimer signaling component^3,7^. Since our antibody of interest, VRC-CH31, was of the same isotype as the VRC01, we kept the sequences and structures of the Fc domains, TM helices, and Igα/Igβ heterodimer as in the 7XQ8^3^ PDB structure and replaced the VRC01 Fab with the VRC-CH31 Fab to model the CH31 BCR using the SWISS-MODELLER^23^ homology modelling webserver. The antigen HIV-1 Env (consisting of gp120) was taken from the 6NNJ^24^ PDB structure and computationally docked into the model CH31 BCR using the HDOCK^25^ webserver to prepare the model CH31 BCR bound by the HIV-1 Env protomer (**Figure 1** and **Supplementary Figure 1**). Two different simulation systems of the model CH31 BCR were prepared in the presence/absence of the antigen (**Figure 1**). A close look at the HIV-1 Env bound Fab domains in the starting model revealed that the HIV-1 Env primarily interacted with the CDR3 regions of one of the Fab domains while it had minor interactions with the other Fab domain as well (**Supplementary Figure 1**). Five different MD simulation replicas of 500 ns each were performed on each CH31 BCR simulation system (**Supplementary Table 1**).

### Increase in the flexibility of BCR upon antigen binding

We calculated the average root-mean-square fluctuations (RMSFs) from all five MD simulation replicas for each BCR simulation system. We then determined the changes in the flexibilities of the BCR upon antigen binding by subtracting the RMSF of the BCR system without the HIV-1 Env from that of the corresponding system bound by the HIV-1 Env (**Figure 2**).

**Figure 2.**
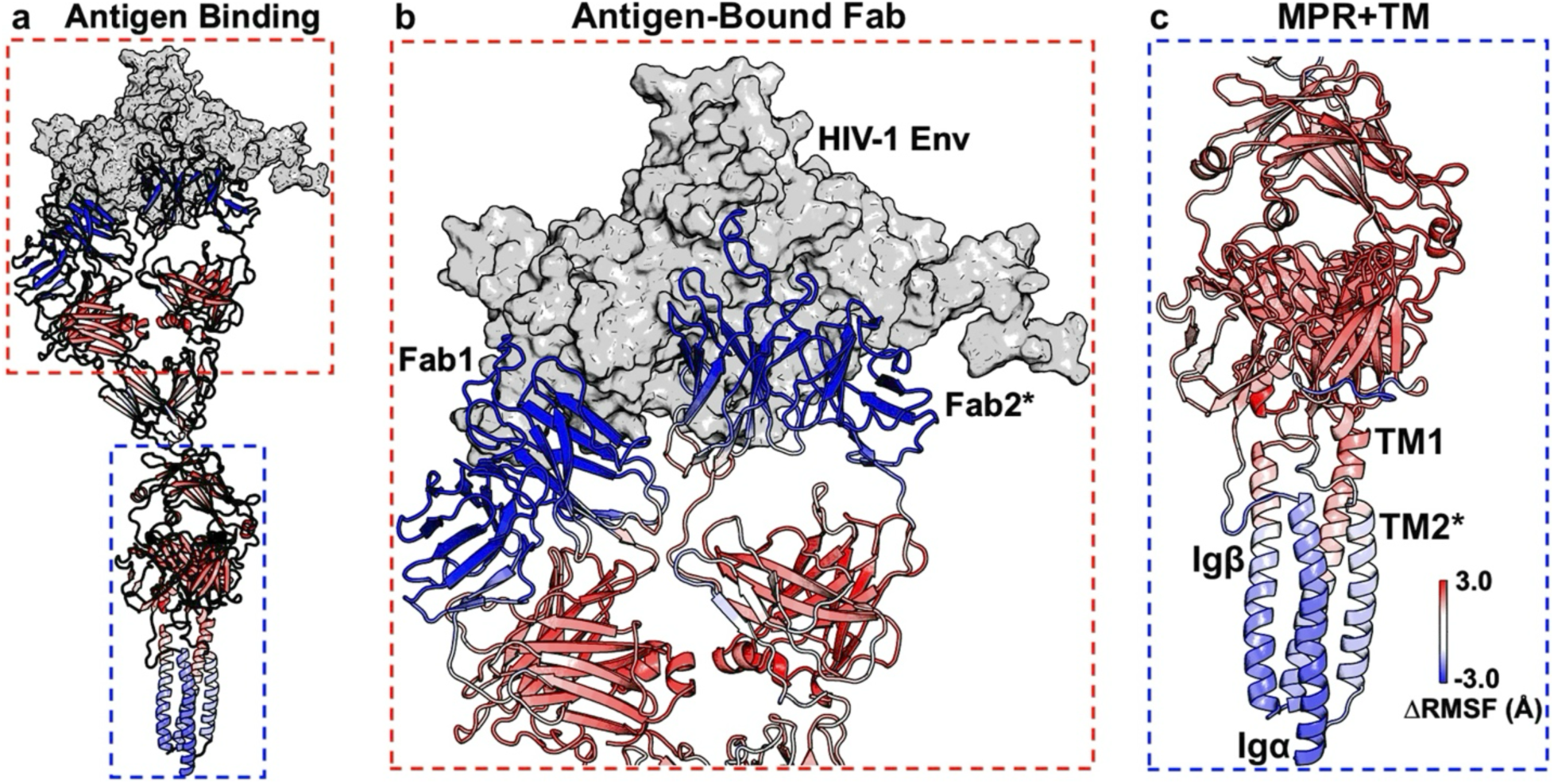
Changes in the system flexibility as measured by the changes in root-mean-square fluctuations (ΔRMSF) upon antigen binding to the BCR. **(a)**. The important domains that showed significant dynamic changes upon antigen binding in the BCR are zoomed in **(b-c)**. A color scale of blue (−3.0) – white (0.0) – red (3.0) is used to show the magnitudes of ΔRMSFs. The HIV-1 Env is colored gray.

Overall, binding of the antigen to the BCR increased the flexibility of multiple domains of the BCR (**Figure 2a**). Antigen binding increased the flexibility of the Fab domains, except for both variable parts (Fv) where the HIV-1 Env bound to the receptor (**Figure 2b**), the whole Fc domains, including their MPR, and the extracellular domains (ECDs) of Igα and Igβ (**Figure 2a** and **2c**). However, the membrane helices turned slightly more rigid upon antigen binding (**Figure 2a** and **2c**). Seemingly, antigen binding to the BCR at the extracellular Fab domains propagated dynamic changes throughout the BCR, opposing the classical cross-linking model of BCR activation^7–10^, and increased the flexibility of the MPR (including the C_H3_ and C_H4_ regions of the Fc domains and ECDs of Igα and Igβ), supporting the conformation-induced oligomerization model^11,12^.

### Allosteric changes in BCR transmembrane helices upon antigen binding

We employed the HELANAL^26^ module within the MDAnalysis^27,28^ python package to calculate the global tilt angles with respect to the vertical axis of the four membrane helices (TM1, TM2, Igα, and Igβ) from the MD simulations of the BCR simulation systems to determine the effects of antigen binding on the orientations of the receptor within the membrane (**Figure 3**). The average and standard deviations of the global tilt angles of the membrane helices calculated from the MD simulations of the BCR systems in the membrane were included in **Figure 3**. The time courses of the global tilt angles were plotted in **Supplementary Figure 2**, and the histograms of the global tilt angles of the four membrane-bound domains sampled from the MD simulations were shown in **Supplementary Figure 3**. Overall, antigen binding in the BCR seemingly shifted the global tilt angles of the membrane helices to lower and narrower distributions (i.e., smaller averages and standard deviations of tilt angles as well as smaller medians and interquartile ranges) (**Figure 3**). Noticeably, the TM2, which belonged to the second IgM subunit that showed the primary interactions with the HIV-1 Env, was less tilted than TM1 (**Figure 3b**-**3c**). The Igα, which interacted more closely with TM2 than TM1, was in turn less tilted than Igβ, which lay closer to TM1 than TM2 (**Figure 3b**-**3c**). In particular, the average global tilt angle of the TM1 shifted from ∼36.1° ± ∼12.2° to ∼27.3° ± ∼6.5°, of the TM2 shifted from 15.1° ± 6.0° to 9.0° ± 4.3°, of the Igα shifted from ∼14.3° ± ∼6.1° to ∼11.2° ± ∼4.9°, and of the Igβ shifted from ∼34.8° ± ∼8.3° to ∼25.5° ± ∼8.8° upon antigen binding (**Figure 3b**). However, given the relatively low number of simulation replicas (*N = 5*) performed for each system, we could not determine whether the shifts were statistically significant based on the unpaired student’s t-test (**Figure 3b**). On the other hand, the boxplots of most global tilt angles of the four membrane-bound domains sampled from the MD simulations also showed clear shifts of the angles towards lower and narrower distributions in the antigen-bound BCR compared to the system without antigen (**Figure 3c**). In particular, the median, Q1, and Q3 of the TM1 tilt angles were found to be ∼37.6°, ∼28.5°, and ∼45.4° for the BCR without antigen and ∼26.7°, ∼22.9°, and ∼31.1° for the BCR with antigen bound (**Figure 3c**). The median, Q1, and Q3 of the TM2 tilt angles were found to be ∼15.3°, ∼8.2°, and ∼21.1° for the BCR without antigen and ∼8.8°, ∼5.9°, and ∼11.8° for the BCR with antigen bound (**Figure 3c**). The median, Q1, and Q3 of the Igα tilt angles were found to be ∼13.6°, ∼9.8°, and ∼18.6° for the BCR without antigen and ∼10.8°, ∼7.7°, and ∼14.3° for the BCR with antigen bound (**Figure 3c**). The median, Q1, and Q3 of the Igβ tilt angles were found to be ∼34.3°, ∼29.2°, and ∼40.4° for the BCR without antigen and ∼24.1°, ∼19.4°, and ∼31.1° for the BCR with antigen bound (**Figure 3c**). Therefore, antigen binding at the extracellular Fab domains reoriented the membrane helices, supporting the conformation-induced oligomerization and dissociation activation models of BCR activation^7,11–15^, and accordingly the enhanced BCR flexibility could disrupt either its self-association^14^ or association with a regulatory co-receptor^29^.

**Figure 3.**
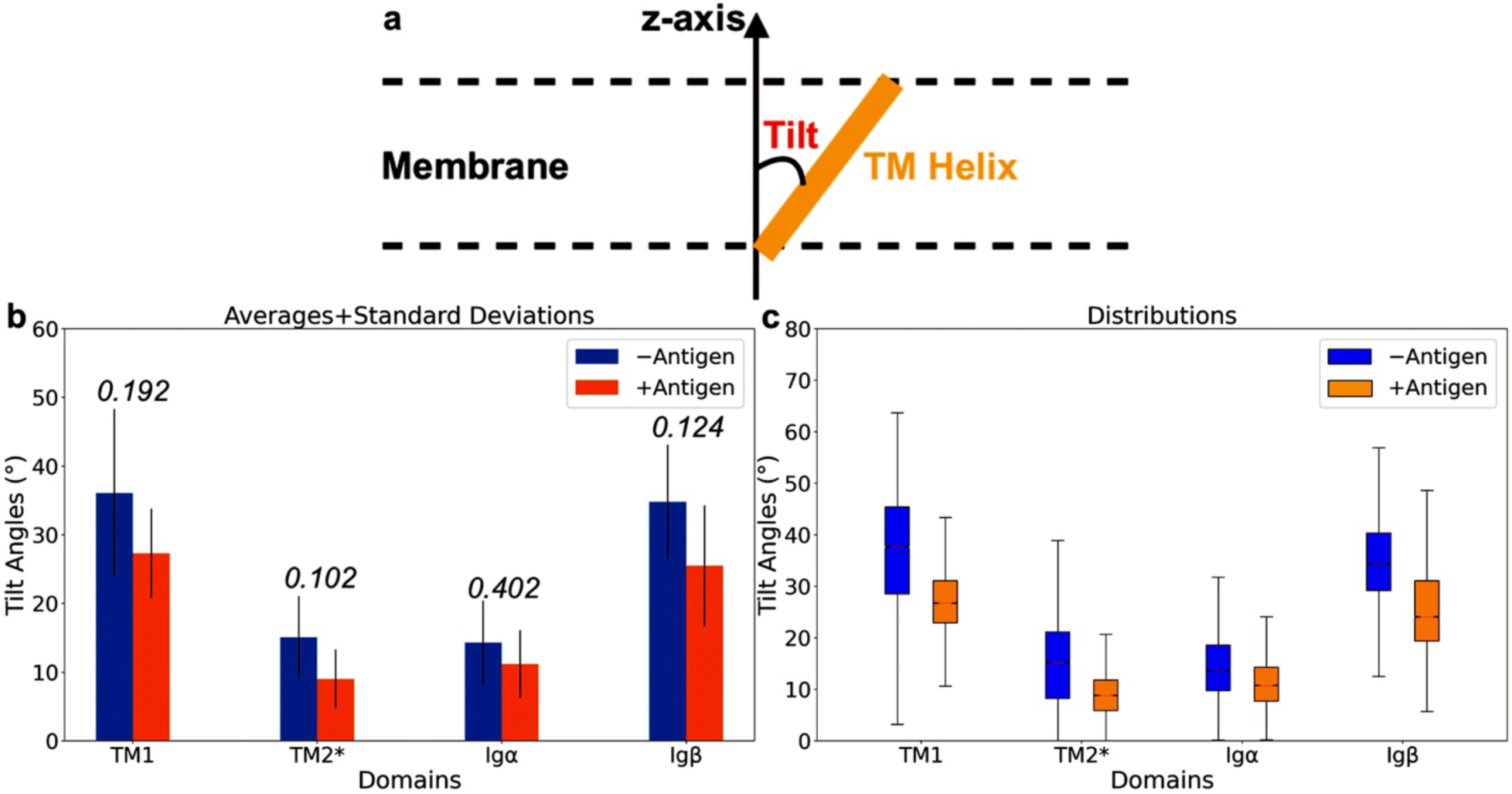
Effects of antigen binding to the BCR on the global tilt angles of the transmembrane helices (including TM1, TM2, and the Igα/Igβ heterodimer) normal to membrane (z-axis) as calculated from the MD simulations of the BCR complexes. **(a)** The global tilt angle was defined as the angle formed by the transmembrane helices and the z- axis (which was perpendicular to the membrane plane). **(b)** The averages and standard deviations of the global tilt angles of the transmembrane helices calculated from the MD simulations of the BCR complexes. The bars for the BCR without and with antigen bound are colored blue and orange, respectively. The p-values were calculated using the unpaired student’s t-test with the numbers of samples being the numbers of simulation replicas (*N = 5*) and shown on top of the bar graphs. **(c)** The distributions of the global tilt angles of the transmembrane helices calculated from the MD simulations of the BCR complexes shown as boxplots. The antigen was found to mostly interact with the second IgM subunit (containing TM2, marked by *).

We then performed principal component analysis (PCA) on the MD simulations of every antigen-bound BCR simulation system to identify the representative low-energy conformational states of the BCR complexes (**Supplementary Figures 4-6**). The BCR was relatively rigid as similar conformations were observed at the Fab domains between different states (**Supplementary Figures 4-6**). Due to the rigidity of the two Fab domains, they always stayed close to each other, causing the model antigen to interact with both Fab domains (**Supplementary Figures 5-6**) that is consistent with the minimal antigen valency requirement for B cell activation^30,31^. Furthermore, as antigen bound to the BCR, the membrane helices turned more upright (**Supplementary Figures 4-5**), being consistent with our calculations of the global tilt angles in **Figure 3**.

### Induction of signaling motifs upon antigen binding

To determine the effects of the antigen binding on the signaling motif within the BCR, we calculated the changes in residue contact frequencies (Δ(frequency)) between Igα and Igβ upon antigen binding in the BCR. The changes in residue contact frequencies between the transmembrane regions of Igα and Igβ upon antigen binding are shown in **Figure 4**. Overall, we observed that the loop that connected the ECD and the membrane helix of Igα wrapped around the Igβ, significantly increased the frequencies of residue contacts within the Igα/Igβ around this region (**Figure 4a**). In particular, notable residue pairs that displayed significantly increased contacts upon antigen binding to the BCR included Igα residue L163 – Igβ residue P176 (with a Δ(frequency) of 0.25), Igα residue E138 – Igβ residue R154 (a possible ionic bond, with a Δ(frequency) of 0.32), Igα residue K141 – Igβ residue K158 (a possible hydrogen bond, with a Δ(frequency) of 0.47), Igα residue N142 – Igβ residue K158 (a possible hydrogen bond, with a Δ(frequency) of 0.51), Igα residue E138 – Igβ residue K158 (a possible ionic bond, with a Δ(frequency) of 0.76), Igα residue F133 – Igβ residue D159 (with a Δ(frequency) of 0.89), Igα residue G137 – Igβ residue N155 (with a Δ(frequency) of 0.93), and Igα residue L134 – Igβ residue D159 (with a Δ(frequency) of 1.0) (**Figure 4a** and **4c**). As residues F133, L134, and G137 of the Igα were all located at the loop connecting the ECD and membrane-bound helix of the Igα, their significant increases in contact with the Igβ confirmed our observation that antigen binding to the BCR led to the wrapping of the Igα connecting loop around Igβ. Furthermore, we observed noticeable reduced contacts towards the C-terminal ends of Igα and Igβ, including Igα residue P159 – Igβ residue V175 (with a Δ(frequency) of −0.25) (**Figure 4a**-**4b**). This result is consistent with the fact that the changes in Igα/Igβ heterodimer signaling component may lead to the exposure of their intracellular ITAM phosphorylation motifs (**Figure 4a**-**4b**) and thereby overcome the autoinhibited state as proposed in the dissociation activation model^13–15,19^.

**Figure 4.**
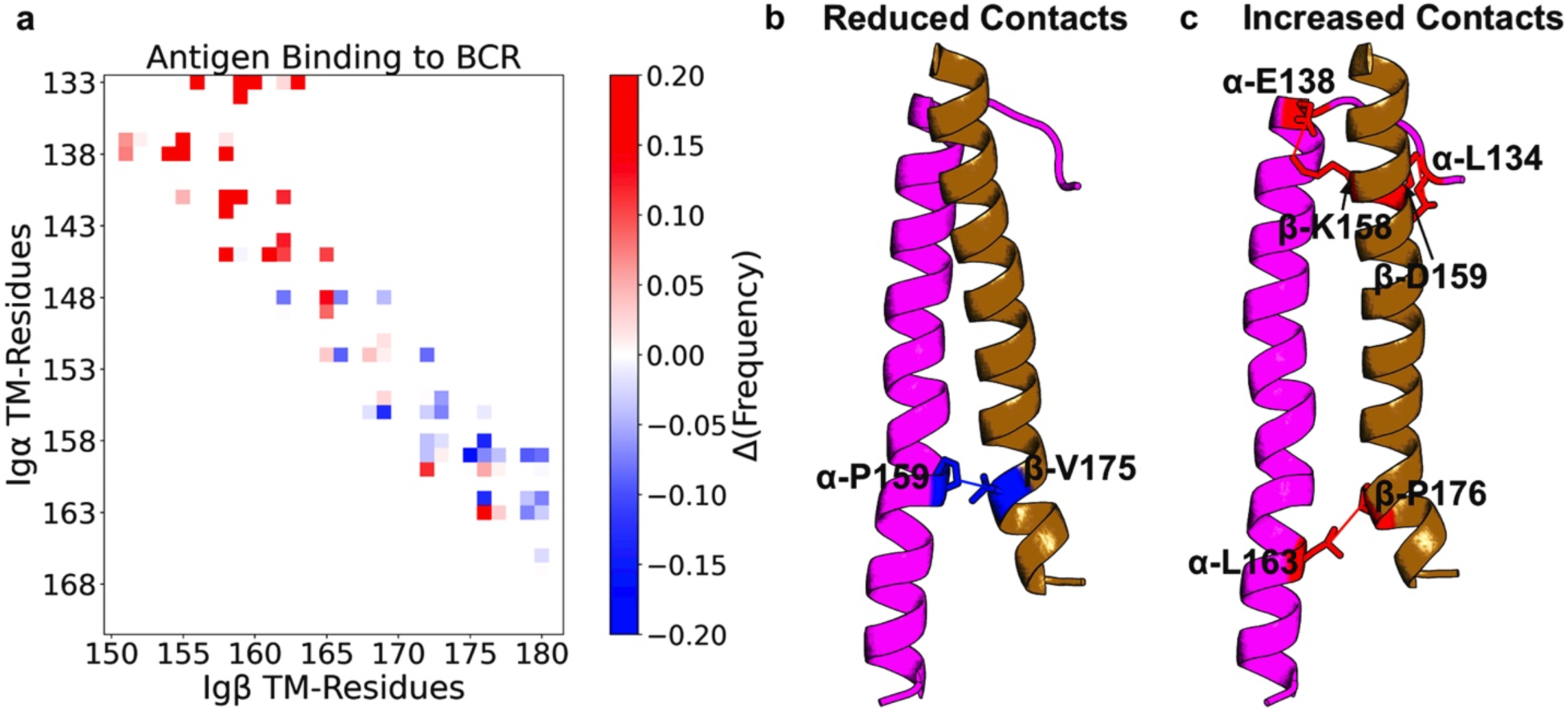
Changes in residue contacts between Igα and Igβ upon binding of the antigen to the membrane bound BCR. **(a)** Changes in residue contact frequencies between Igα and Igβ upon HIV-1 Env binding to the BCR. A contact definition of ≤ 8.0 Å distance between Cα atoms was used. A color scale of blue (−0.2) – white (0) – red (0.2) was used to show the magnitudes of changes in residue contact frequencies. **(b)** Representative reduced contacts upon antigen binding in the BCR. **(c)** Representative increased contacts upon antigen binding in the BCR. The structural residue contacts colored blue were those that showed decreases in contact frequencies upon antigen binding, whereas those colored red showed increased in contact frequencies upon antigen binding.

### Lipid reorganization in response to antigen-induced conformational changes in BCR

In addition to the AA MD simulations, we also performed long timescale coarse grained (CG) molecular dynamics simulations for 5µs, started from the final conformations of each AA MD simulation replica obtained from each BCR simulation system to determine lipid reorganization upon antigen binding in the BCR (**Supplementary Figure 7**). We counted the number of lipid molecules within 4Å of the BCR within the last 100 ns of the AA and CG MD simulations of the CH31 BCR simulation systems using the MDAnalysis^27,28^ python package (**Figure 5a**).

**Figure 5.**
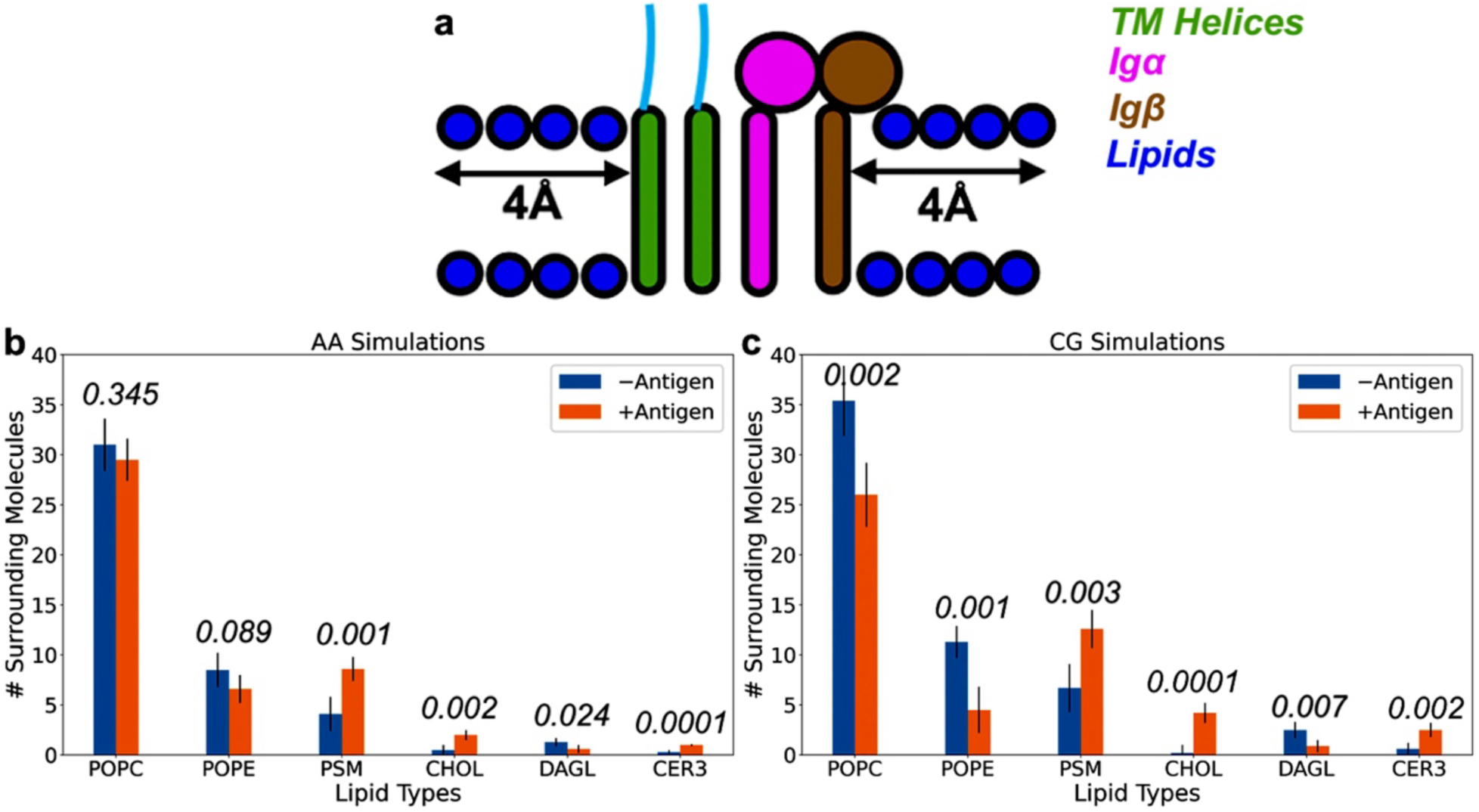
Reorganization of the local lipid environments upon antigen binding to the BCR in the complex membrane. **(a)** All lipid molecules within 4Å distance of the membrane-peripheral region and membrane-bound helices of the BCR are considered. **(b-c)** The numbers of surrounding lipid molecules during the last 100 ns of the AA **(b)** and CG MD simulations **(c)** in the complex membrane are shown. The bars for the BCR without and with antigen bound are colored blue and orange, respectively. The p-values were calculated using the unpaired student’s t-test with the numbers of samples being the numbers of simulation replicas (*N = 5*) and shown on top of the bar graphs.

The initial AA conformation of the CH31 BCR was surrounded by 35 POPC, 8 POPE, 5 PSM, 2 CHOL, 1 DAGL, and 0 CER3 lipid molecules (**Supplementary Figure 8**). On average, 31.0 ± 2.6 POPC, 8.5 ± 1.7 POPE, 4.1 ± 1.7 PSM, 0.5 ± 0.5 CHOL, 1.3 ± 0.4 DAGL, and 0.3 ± 0.2 CER3 lipid molecules came into contacts with the BCR within the last 100ns of its five AA MD simulations (**Figure 5b**). At the end of the five CG MD simulations, an average number of 35.4 ± 3.5 POPC, 11.3 ± 1.6 POPE, 6.7 ± 2.4 PSM, 0.2 ± 0.8 CHOL, 2.5 ± 0.8 DAGL, and 0.6 ± 0.6 CER3 lipid molecules contacted the receptor (**Figure 5c**). Meanwhile, at the start of the AA MD simulation of the antigen bound BCR, 35 POPC, 7 POPE, 8 PSM, 2 CHOL, 1 DAGL, and 0 CER3 lipid molecules were in contact with the BCR (**Supplementary Figure 8**). In the last 100ns of the five AA MD simulations of the system, 29.5 ± 2.1 POPC, 6.6 ± 1.4 POPE, 8.6 ± 1.2 PSM, 2.0 ± 0.5 CHOL, 0.6 ± 0.4 DAGL, and 1.0 + 0.1 CER3 lipid molecules surrounded the receptor (**Figure 5b**). At the end of the five CG MD simulations, on average 26.0 ± 3.2 POPC, 4.5 ± 2.3 POPE, 12.6 ± 1.9 PSM, 4.2 ± 1.0 CHOL, 0.9 ± 0.6 DAGL, and 2.5 ± 0.7 CER3 were within 4Å distance of the receptor (**Figure 5c**).

We observed noticeable increases in the numbers of ordered lipid molecules such as PSM, CHOL, and CER3 in the proximity of the BCR upon antigen binding while seeing significant decreases in the numbers of disordered lipid molecules, including POPC, POPE, and DAGL surrounding the BCR (**Figure 5b**-**5c**). The reorganizations of lipid molecules surrounding the BCR upon antigen binding were determined to be mostly statistically significantly during the AA MD simulations, except for the POPC and POPE lipid molecules, (**Figure 5b**) and all statistically significant during the CG MD simulations (**Figure 5c**). Therefore, conformational changes that occurred due to the binding of the antigen led to lipid reorganization within the proximity of the BCR as the average numbers of different lipid molecules surrounding the BCR were different. This raises the possibility that activated BCR may prefer a different lipid composition than those that are in basal state, being consistent with previous studies^7,11–15^. It is notable that on naïve B cells, the IgM class BCRs are clustered in distinct protein/lipid islands and following antigen binding co-localizes with CD19 and lipid ordered domains^32^ and thus consistent with our results showing an increase in ordered lipid molecules.

### Membrane lipid properties during the MD simulations of the BCR complexes

We used the LiPyphilic^33^ python toolkit to calculate the area per lipid in different membrane environments for both the AA and CG simulations. The average area per lipid molecule was 58.0 Å^2^ in the BCR simulation system, while the area per lipid molecule was 57.6 Å^2^ in the antigen-bound BCR simulation system. During the CG MD simulations, the average areas per lipid molecule for the BCR and HIV-1 Env-bound BCR were 57.5 Å^2^ and 57.2 Å^2^, respectively. Seemingly, the average areas per lipid were comparable between the AA and CG MD simulations. Furthermore, antigen binding to the BCR appeared to slightly increase the order of membrane lipids in the bilayers as we saw small decreases in the average areas per lipid upon antigen binding.

To confirm our predictions regarding the membrane orders, we further employed the LiPyphilic^33^ python toolkit to calculate the average membrane thicknesses obtained from the AA and CG MD simulations of the BCR in the presence/absence of the model HIV-1 Env antigen (**Supplementary Table 2**). In particular, the average membrane thickness calculated from the AA simulations increased from 36.5 ± 1.8 Å for the BCR to 42.2 ± 4.2 Å for the HIV-1 Env bound BCR (**Supplementary Table 2**). The average membrane thickness calculated from the CG simulations was comparable to the AA simulations, with the respective values being 38.1 ± 6.1 Å and 48.8 ± 8.2 Å for the BCR and HIV-1 Env bound BCR (**Supplementary Table 2**). Seemingly, upon antigen binding to the BCR, the average membrane thickness increased in a statistically significant manner (**Supplementary Table 2**), being consistent with our finding that antigen-bound BCR preferred ordered to disordered lipid bilayers in **Figure 5**.

We also employed the *2D streamplots*^34^ module within the *MDAnalysis*^27,28^ simulation analysis toolkit to monitor the lipid diffusions during our AA and CG MD simulations (**Supplementary Figures 9-12**). Overall, we observed higher lipid diffusion speeds during the CG MD simulations than the AA MD simulations. In particular, the maximum speeds observed for lipid diffusions during the AA MD simulations of the BCR and HIV-1 Env-bound BCR were between ∼2.6 - ∼2.7 (**Supplementary Figures 9-10**), while the maximum speeds observed for lipid diffusions during the CG MD simulations were ∼4.0-4.2 (**Supplementary Figures 11-12**). We then used the LiPyphilic^33^ python toolkit to calculate the average lipid diffusion coefficients within the membranes of the different AA and CG BCR simulation systems (**Supplementary Table 3**). On average, the CG lipid diffusion coefficients were significantly higher than the AA lipid diffusion coefficients, illustrating by the p-values from the unpaired student’s t-test. In particular, the pooled diffusion coefficients calculated from the AA and CG MD simulations of the BCR were 1.7×10^-6^ ± 2.7×10^-7^ nm^2^/ns and 1.2×10^-5^ ± 5.1×10^-6^ nm^2^/ns, and of the HIV-1 Env bound BCR were 1.4×10^-6^ ± 2.2×10^-7^ nm^2^/ns and 9.8×10^-6^ ± 3.3×10^-6^ nm^2^/ns (**Supplementary Table 3**). On the other hand, we found no significant difference in the lipid diffusion coefficients in the presence and absence of the antigen in the BCR (**Supplementary Table 3**). Therefore, the CG MD simulations facilitated lipid motions within the membranes compared to the AA MD simulations due to the flattened free energy surfaces as a result of the employed MARTINI 3^35^ force field during the CG simulations. However, it should be mentioned that the flattened free energy surfaces created by the CG simulations with MARTINI 3^35^ force field yield at least four times higher diffusion coefficients than those calculated from all-atom MD simulations. Rather, our intention here was to show the lipid molecules diffused significantly faster during CG simulations using MARTINI 3^35^ force field than AA simulations^36–38^ and hence justified our usage of CG simulations for accelerating lipid reorganization.

## Discussion

Recently, the first cryo-EM structure of the human BCR of the IgM isotype was published^3^. Given the new structural information of this asymmetric complex, we have carried out extensive MD simulations on different complexes of the model CH31 BCR. We profiled the important protein conformations and protein-lipid interactions upon antigen (HIV-1 envelope protein) binding in a complex membrane (**Figure 1**). We related the resulting dynamics captured from the simulations to known models of B-cell activation.

Findings from our work support either the conformation-induced oligomerization model^11,12^ or the dissociation activation model^7,13–15^ while opposing the classical cross-linking model^7–10^ of BCR activation. The classical cross-linking model of BCR activation states that no signal propagation from the extracellular domain to the cytoplasm was necessary for BCR activation^7–10^. Instead, BCR activation is a simple event of the Igα/Igβ aggregation accomplished by binding multivalent antigens to multiple BCRs^7–10^. This work shows that antigen binding at the extracellular Fab domains led to changes in the dynamics throughout the BCR. This is required for the conformation-induced oligomerization and dissociation activation models (i.e., the conformational-change induced activation models). Furthermore, the flexibility of the BCR monomer and the lipid reorganization around the BCR upon antigen binding favors dissociation activation model even though our simulations were performed on a single BCR complex. It should be noted that we do not rule out the fact that multiple activation mechanisms can coexist.

We identify three critical BCR conformational changes upon antigen binding that show the pathway to its activation. First, antigen binding increased the flexibility of the MPR (including the C_H3_ and C_H4_ regions of the Fc domains and the ECDs of Igα/Igβ) (**Figure 2**), being consistent with the conformation-induced oligomerization model^11,12^ as the model proposed that antigen binding opened an oligomerization interface at the MPR. Second, antigen binding increased residue contacts in the transmembrane helices near the MPR of Igα and Igβ, while reduced residue contacts of the Igα and Igβ in the inner membrane leaflet, being consistent with the dissociation activation model^7,13–15^ (**Figure 4**). Third, antigen binding in the BCR consistently shifted the global tilt angles of the transmembrane helices (including TM1, TM2, Igα, and Igβ) of the BCR to lower and narrower distributions as shown by the boxplots of tilt angle distributions (**Figures 3**). The changes in the tilting of the transmembrane helices of Igα/Igβ heterodimer signaling component can alter the exposure of the ITAM motifs in the cytoplasmic tails required for the downstream signaling. In the mouse IgM BCR structure^19^, an autoinhibition model was proposed involving a helical ITAM motif fold-back onto Igβ TMD and therefore, suggesting that antigen binding could induce ITAM exposure for phosphorylation. We find the dynamical motions governing these events are correlated. However, we cannot state the order of events from the computational approaches taken here.

Lipid reorganization around BCR upon antigen-induced conformational changes indirectly supports the dissociation activation model and the class-specific organization of BCRs on naïve B cells. We observe that the antigen-bound BCR preferred to be surrounded by more ordered lipid molecules such as PSM, CHOL, and CER3, while the free BCR is surrounded by more disordered lipid molecules such as POPC, POPE, and DAGL (**Figure 5**). These lipid rearrangements are caused by changes in the MPR and transmembrane regions of BCR because of antigen binding. This indicates that, potentially, the antigen-bound BCR prefers a different lipid environment than the free one. Specifically, the shifts in tilting towards the normal of the membrane may prefer a composition that is a more ordered domain with a higher membrane thickness. This could be potentially driven by the hydrophobic mismatch caused by the transmembrane rearrangements (**Figures 2**-**4**). These observations are supported by previous studies that suggested a relocation of BCR from a disordered domain to an ordered domain^7,11–15^.

In conclusion, we have uncovered three critical dynamical events that could be associated with antigen-dependent activation of BCR. First, antigen binding caused increased flexibility in regions distal to the antigen binding site. Second, antigen binding resulted in alterations of IgM transmembrane helices and MPR regions. Third, these alterations, impacted the relative interaction between Igα and Igβ and the orientations of their transmembrane helices. These changes are expected to influence the exposure or phosphorylation of ITAM motifs in the cytoplasmic tails of Igα/Igβ. These conformational changes could potentially relocate BCR in a different region of the membrane as indicated by the differential preferences of lipids before and after the antigen binding. Even though the simulations considered only a single BCR complex, our work indirectly supports the conformational-change induced activation models. Further simulation studies involving multiple BCRs in different membrane environments performed for much longer timescales are warranted to make the direct connection to the proposed activation models conclusively.

## Materials and Methods

### Simulation system setups

We started from the cryo-EM structure of human B-cell antigen receptor (BCR) of the IgM isotype with the Fab domains of the VRC01 antibody (PDB: 7XQ8)^3^. Since our antibody of interest VRC-CH31 was of the same isotype as the VRC01 antibody, the sequences and structures of the Fc domains, transmembrane helices, Igα, and Igβ domains in the BCR with the Fab domains of VRC-CH31 should be identical to those from the 7XQ8 PDB^3^ structure. The sequences of the VRC-CH31 Fab heavy and light chains were retrieved from the UniProt database^39^ with UniRef IDs of UPI00038A5F4F and UPI00038A5F3B, respectively, and attached to the rest of the BCR through sequence alignment and SWISS-MODEL template-based homology modelling webserver^23^. SWISS-MODEL did not model the intracellular loops of the Igα/Igβ heterodimer^3^ even when we provided the full sequences from the 7XQ8 FASTA file (**Supplementary Figure 13a**). We also tried using AlphaFold2-Multimer^40,41^ and AlphaFold3^42^ in modelling the Igα/Igβ heterodimer with intracellular loops as well as the full BCR structure (**Supplementary Figure 13b-13d**). We observed that AlphaFold^40–42^ predicted the Igα/Igβ heterodimer intracellular loop to “U-turn” into the membrane (**Supplementary Figure 13b-13d**). Therefore, we decided to use the BCR model built by SWISS-MODELLER^23^ to proceed with our simulations. The sequences and structure of the HIV-1 envelope protein (including the gp120 envelope protein) were taken from the 6NNJ PDB structure^24^ and remodeled to fill in the missing regions using the SWISS-MODEL webserver^23^. The HIV-1 envelope protein was docked into the CH31 BCR using the HDOCK integrated protein-protein docking webserver^25^. To examine both the effects of antigen binding on the dynamics mechanisms of the BCR with VRC-CH31 Fab domains, we set up two different simulation systems, including the CH31 BCR and CH31 BCR bound by the HIV-1 envelope protein (HIV- 1 Env) in the complex membrane (**Figure 1**). The CHARMM-GUI webserver^43–46^ was used to set up the initial atomistic simulation systems. We did not model the glycosylation of the CH31 BCR and the HIV-1 envelope protein. The CH31 BCR systems were embedded in the membrane lipid bilayer before being solvated in 0.15 M NaCl solutions. The resulting system sizes ranged from ∼1.5 million (for the BCR without HIV-1 Env) to ∼2.5 million atoms (for the BCR bound by HIV-1 Env). The CHARMM36m force field parameter sets^47^ were used for the proteins and lipids in the all-atom (AA) molecular dynamics (MD) simulations. The elastic networks of MARTINI 3 force field parameter sets^35^ were used in the coarse-grained (CG) MD simulations.

### Multiscale molecular dynamics simulation protocols

In the multiscale MD simulations, AA simulations were iterated with long timescale CG simulations (**Supplementary Figure 7**). The protein-protein interactions were refined with AA simulations, while lipid mixing was facilitated during the CG simulations. Here, however, CG simulations were only performed to examine the lipid mixing properties during the AA simulations. In the AA MD simulations, periodic boundary conditions were applied to the simulation systems, and bonds containing hydrogen atoms were restrained with the LINCS^48^ algorithm. The electrostatic interactions were calculated using the particle mesh Ewald (PME) summation^49^ and the Verlet cutoff scheme^50^ with a cutoff distance of 12 Å for long-range interactions. The temperature was kept constant at 310 K using the Nose-Hoover thermostat^51^ with a friction coefficient of 1.0 ps^-1^. The pressure was kept constant at 1.0 bar using the Parrinello-Rahman barostat^52^ with semi-isotropic coupling. The pressure coupling constant was set to 5 ps, and the compressibility was set to 4.5×10^-5^ bar^-1^. The simulation systems were energetically minimized to a maximum of 5,000 steps using the steepest-descent algorithm. Position restraints were applied on the backbone atoms with a force constant of 4,000 kJ.mol^-1^.nm^-2^, on the side chain atoms with a force constant of 2,000 kJ.mol^-1^.nm^-2^, and on the lipids and dihedral angles with a force constant of 1,000 kJ.mol^- 1^.nm^-2^. The systems were then equilibrated with the constant number, volume, and temperature (NVT) ensemble for a total of 375,000 steps, with a time step of 1 fs used. The force constants for position restraints were gradually reduced from 4,000 to 2,000 to 1,000 kJ.mol^-1^.nm^-2^ for backbone atoms, from 2,000 to 1,000 to 500 kJ.mol^-1^.nm^-2^ for side chain atoms, from 1,000 to 400 kJ.mol^-1^.nm^-2^ for lipids, and from 1,000 to 400 to 200 kJ.mol^-1^.nm^-2^ for dihedral angles, after every 125,000 steps. The systems were further equilibrated with the constant number, pressure, and temperature (NPT) ensemble for a total of 750,000 steps, with a time step of 2 fs used. The force constants for position restraints were gradually reduced from 500 to 200 to 50 kJ.mol^-1^.nm^-2^ for backbone atoms, from 200 to 50 to 0 kJ.mol^- 1^.nm^-2^ for side chain atoms, from 200 to 40 to 0 kJ.mol^-1^.nm^-2^ for lipids, and from 200 to 100 to 0 kJ.mol^-1^.nm^-2^ for dihedral angles, after every 250,000 steps. Finally, the systems were equilibrated with a short 25ns conventional MD (cMD) simulation using a time step of 2 fs. Five 500ns cMD production simulations were then performed on each of the BCR simulation systems. The final frame from each AA MD simulation replica of every BCR simulation system was extracted and stripped oW all ions and water molecules. The protein and membrane lipid portions were converted to their corresponding CG representation using the martinize2 and vermouth framework (https://github.com/marrink-lab/vermouth-martinize)^53^ for the protein, and the *backward.py* (https://github.com/Tsjerk/MartiniTools/blob/master/backward.py) script^54^ for the membrane lipids. The CG models of the proteins were built with the side chain corrections (-scfix option)^55^ and the elastic network (-elastic option)^56^. The elastic bond force constant was set to 700 kJ.mol^-1^.nm^-2^ (-ef 700), the lower and upper bound of the elastic bond cutoff were set to 0.5 and 0.9 nm, respectively (-el 0.5 -eu 0.9)^56^. The elastic bond decay factor and power were both set to 0 (-ea 0 -ep 0) to make the bond strengths independent of bond length^56^. The *Insane* (https://github.com/Tsjerk/Insane) software package^57^ was used to re-solvate the CG systems of the BCR assemblies and lipids in 0.15 M NaCl solutions, with the box dimensions kept identical to the AA simulations. In the CG MD simulations, periodic boundary conditions were applied to the simulation systems. A time step of 20 fs was used. The electrostatic interactions were calculated using the reaction field method^58^ and the Verlet cutoff scheme^50^ with a cutoff distance of 11 Å for long-range interactions. The temperature was kept constant at 310 K using the velocity rescaling thermostat^59^ with a friction coefficient of 1.0 ps^-1^. The pressure was kept constant at 1.0 bar first using the stochastic cell rescaling (C-rescale)^60^ during the equilibration stage and then Parrinello-Rahman barostat^52^ during the production simulations with semi-isotropic coupling. The pressure coupling constant was set to 5 ps, and the compressibility was set to 3×10^-4^ bar^-1^. The simulation systems were energetically minimized to a maximum of 500,000 steps given the large simulation system sizes. They were then equilibrated with the constant number, pressure, temperature (NPT) ensemble for a total of 300,000 steps. Position restraints were applied on the systems, with a force constant gradually reduced by half starting from 1,000 to 50 kJ.mol^-1^.nm^-2^ after every 50,000 steps, except the lipid heads where the force constants were gradually reduced from 200 to 10 kJ.mol^-1^.nm^-2^. The simulation systems were further equilibrated with a short 100ns cMD simulation. Finally, one 5μs cMD production simulation was performed on each CG simulation system obtained from each replica of the previous AA simulations. All simulations were carried out using the *gmx grompp and gmx mdrun* commands in GROMACS 2022^61^ on Los Alamos National Laboratory (LANL) High Performance Computing (HPC) Clusters (**Supplementary Table 1**).

### Simulation analysis

First, we used both the CPPTRAJ^62^ simulation analysis tool and the GROMACS 2022^61^ simulation package to calculate the changes in root-mean-square fluctuations (ΔRMSF) to determine the changes in flexibility in different domains of the BCR complexes upon the binding of the HIV-1 Env to the BCR. Second, we used the HELANAL^26^ module within the MDAnalysis^27,28^ python package to calculate the global tilt angles with respect to the vertical axis of the four membrane helices (TM1, TM2, and the Igα/Igβ heterodimer) of the CH31 BCR simulation systems. Third, we performed the principal component analysis (PCA) to determine the important low-energy conformational states observed for each BCR simulation system. Fourth, we used the native contacts module within the MDAnalysis^27,28^ python package to calculate the changes in residue contact frequencies between the Igα and Igβ upon antigen binding. The native contacts module^27,28^ was also used to calculate the number of lipid molecules within 4Å distance of the BCR within the last 100ns of the simulation systems. Fifth, we used the LiPyphilic^33^ python toolkit to calculate the average area per lipid, average membrane thickness, and average lipid diffusion coefficients in each BCR simulation system. We employed the unpaired student’s t-test (https://www.graphpad.com/quickcalcs/ttest1/?format=SD) to examine the statistical significance of the differences between the BCR simulation systems with and without antigens bound as well as between their AA and CG simulations. Considering configurations from simulations separated by as high as 20 ns as independent samples, we obtained a very low p-value of (i.e., p ≤ 0.0001). However, we were concerned about using independent sampling assumption within a single simulation as changes in conformations at previous timepoints would cause the changes in conformations in the current and later timepoints. Therefore, we used the number of simulation replicas for each system (N = 5) as the number of samples for each system for the p-value calculation. We believed that the outcome of one simulation replica did not affect the outcome of another, so our samples satisfied the independence assumption for the unpaired student’s t-test. However, it should be noted that the unpaired student’s t-test might not be able to detect the statistical significance, given the small number of samples (replicas). Typically, one needs to include much longer simulations and more simulation replicas, depending on the time scales of dynamical processes under consideration, to reliably determine the statistical significance of dynamical processes in MD simulations. Finally, we employed the *2D streamplots*^34^ module within the *MDAnalysis*^27,28^ simulation analysis toolkit to monitor the lipid diffusion speed during the AA and CG MD simulations.

## Data Availability

Data supporting the findings of this study are included in the article and its Supplementary Information files.

## Code availability

This study utilized the standard builds of the simulation software GROMACS 2022^61^ (https://manual.gromacs.org/2022/index.html) for running AA and CG MD simulations with all parameters specified in the Methods section. The related tools used for simulations were specified in the Methods section.

## Acknowledgements

We would like to thank Cesar Lopez, Tyler Reddy, and Nick Hengartner for providing us with valuable insights and suggestions. This study was supported by the National Institute of Allergy and Infectious Diseases of the National Institutes of Health and by the Duke Center for HIV Structural Biology, grant number U54-AI170752-01. The authors would also like to thank the computational resources provided by LANL Institutional Computing (account W23_DCHSB). This work was performed at the Los Alamos National Laboratory, which is operated by Triad National Security, LLC, for the National Nuclear Security Administration of the U.S. Department of Energy (contract 89233218CNA000001). This work is approved for public release and unlimited distribution under LA-UR-24-26941.

## Author Contributions

H.N.D. performed research, analyzed data, and wrote the manuscript. M.Z. contributed to the writing of the manuscript. S.M.A. conceived the study and contributed to the writing of the manuscript. S.G. conceived the study, supervised the research, analyzed data, and wrote the manuscript. All authors contributed towards the final version of the manuscript.

## Competing Interest Declaration

The authors declare no competing interests.

## Notes

### Competing Interest Statement

The authors have declared no competing interest.

### Summary of Updates

Main updates includes the revision of Figure 3 and the addition of statistical tests to calculate p-values and determine the significances in the differences between the properties of B-cell receptors in the presence and absence of an antigen bound. Associated texts were updated accordingly to accompany the new figure and results.

