## Supplementary Information for "Dynamics and Activation of Membrane-Bound B Cell Receptor Assembly"

**Supplementary Table 1. Summary of the MD simulations of the model CH31 BCR in the presence/absence of the model protomer HIV-1 envelope protein (HIV-1 Env) antigen.** Here, AA_MD stands for all-atom molecular dynamics simulations; and CG_MD stands for coarse-grained molecular dynamics simulations.

| ***Simulation System*** | ***Simulation Method*** | ***Simulation Length*** |
|---|---|---|
| CH31 BCR | AA_MD | 5 x 500ns |
| | CG_MD | 5 x 5000ns |
| CH31 BCR bound by monomer HIV-1 Env | AA_MD | 5 x 500ns |
| | CG_MD | 5 x 5000ns |

**Supplementary Table 2. Membrane thickness (Å) calculated from the all-atom (AA) and coarse-grained (CG) MD simulations of the CH31 BCR with and without the model HIV-1 Env antigen bound.** The p-values were calculated using the unpaired student's t-test with the numbers of samples being the numbers of simulation replicas ($N = 5$).

| | ***-Antigen*** | ***+Antigen*** | ***p-values*** |
|---|---|---|---|
| ***AA MD*** | 36.5 ± 1.8 Å | 42.2 ± 4.2 Å | 0.024 |
| ***CG MD*** | 38.1 ± 6.1 Å | 48.8 ± 8.2 Å | 0.047 |

**Supplementary Table 3. Diffusion coefficients ($nm^2/ns$) of the lipid molecules calculated from the AA and CG MD simulations of the CH31 BCR with and without the model HIV-1 Env antigen bound.** The p-values were calculated using the pooled average and standard deviation diffusion coefficients obtained from the five (*N* = *5*) simulation replicas per simulation system per AA or CG representation.

| | ***-Antigen*** | ***+Antigen*** | |
|---|---|---|---|
| ***AA Sim1*** | $1.6 \times 10^{-6} \pm 7.6 \times 10^{-8}$ | $1.5 \times 10^{-6} \pm 8.2 \times 10^{-8}$ | |
| ***AA Sim2*** | $1.5 \times 10^{-6} \pm 7.1 \times 10^{-8}$ | $1.2 \times 10^{-6} \pm 7.9 \times 10^{-8}$ | |
| ***AA Sim3*** | $2.2 \times 10^{-6} \pm 4.7 \times 10^{-8}$ | $1.2 \times 10^{-6} \pm 8.0 \times 10^{-8}$ | |
| ***AA Sim4*** | $1.5 \times 10^{-6} \pm 6.4 \times 10^{-8}$ | $1.2 \times 10^{-6} \pm 1.0 \times 10^{-7}$ | |
| ***AA Sim5*** | $1.6 \times 10^{-6} \pm 7.0 \times 10^{-8}$ | $1.7 \times 10^{-6} \pm 1.0 \times 10^{-7}$ | |
| ***CG Sim1*** | $4.3 \times 10^{-6} \pm 3.5 \times 10^{-7}$ | $6.9 \times 10^{-6} \pm 5.6 \times 10^{-7}$ | |
| ***CG Sim2*** | $8.9 \times 10^{-6} \pm 5.2 \times 10^{-7}$ | $9.7 \times 10^{-6} \pm 7.4 \times 10^{-7}$ | |
| ***CG Sim3*** | $1.9 \times 10^{-5} \pm 4.6 \times 10^{-7}$ | $8.0 \times 10^{-6} \pm 6.9 \times 10^{-7}$ | |
| ***CG Sim4*** | $1.5 \times 10^{-5} \pm 2.7 \times 10^{-7}$ | $8.5 \times 10^{-6} \pm 6.5 \times 10^{-7}$ | |
| ***CG Sim5*** | $1.1 \times 10^{-5} \pm 6.2 \times 10^{-7}$ | $1.6 \times 10^{-5} \pm 8.9 \times 10^{-7}$ | |
| | ***-Antigen*** | ***+Antigen*** | ***p-values*** |
| ***Pooled AA*** | $1.7 \times 10^{-6} \pm 2.7 \times 10^{-7}$ | $1.4 \times 10^{-6} \pm 2.2 \times 10^{-7}$ | 0.077 |
| ***Pooled CG*** | $1.2 \times 10^{-5} \pm 5.1 \times 10^{-6}$ | $9.8 \times 10^{-6} \pm 3.3 \times 10^{-6}$ | 0.52 |
| ***p-values*** | 0.002 | 0.0004 | |

**Supplementary Figure 1. Starting conformation of the CH31 BCR bound by the model HIV-1 envelope protein (HIV-1 Env) antigen.** The model antigen stayed close to both Fab domains in the starting model, interacting mostly with the second IgM subunit (containing Fab2). The Fab heavy chains are colored red, Fab light chains are colored blue, Fc domains are colored cyan, transmembrane helices of BCR are colored green, Igα is colored magenta, Igβ is colored brown, and the HIV-1 Env is colored orange.

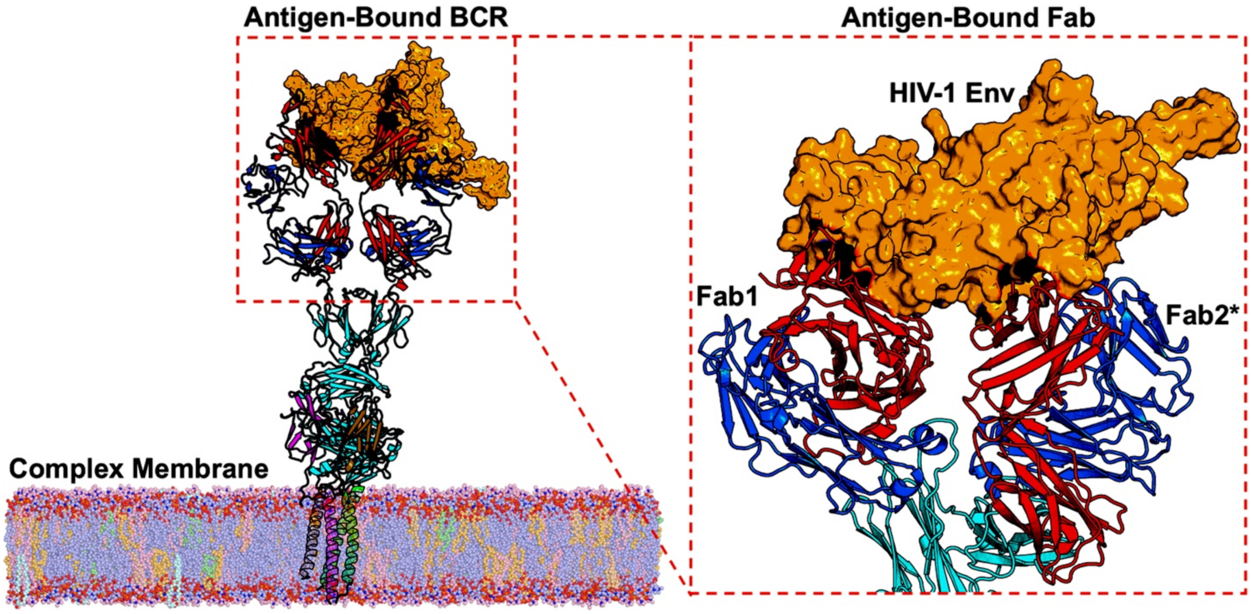

**Supplementary Figure 2. Time courses of the global tilt angles of the transmembrane helices (including TM1, TM2, and the Igα/Igβ heterodimer) with respect to the vertical axis calculated from the MD simulations of the BCR complexes in the presence/absence of the model HIV-1 Env antigen. (a)** Time courses of the global tilt angles of the TM1, TM2, Igα, and Igβ calculated from the five simulation replicas of the BCR. **(b)** Time courses of the global tilt angles of the TM1, TM2, Igα, and Igβ calculated from the five simulation replicas of the BCR bound by HIV-1 Env.

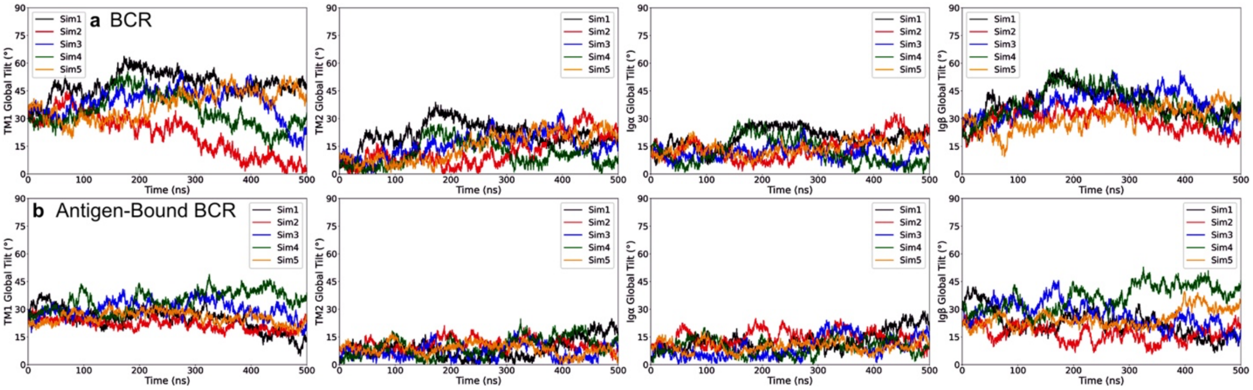

**Supplementary Figure 3. Histograms of the global tilt angles of the transmembrane helices (including TM1, TM2, and the Igα/Igβ heterodimer) with respect to the vertical axis calculated from the MD simulations of the BCR complexes in the presence/absence of the model HIV-1 Env antigen. (a)** Distributions of the global tilt angles of the TM1, TM2, Igα, and Igβ calculated from the five simulation replicas of the BCR. **(b)** Distributions of the global tilt angles of the TM1, TM2, Igα, and Igβ calculated from the five simulation replicas of the BCR bound by HIV-1 Env.

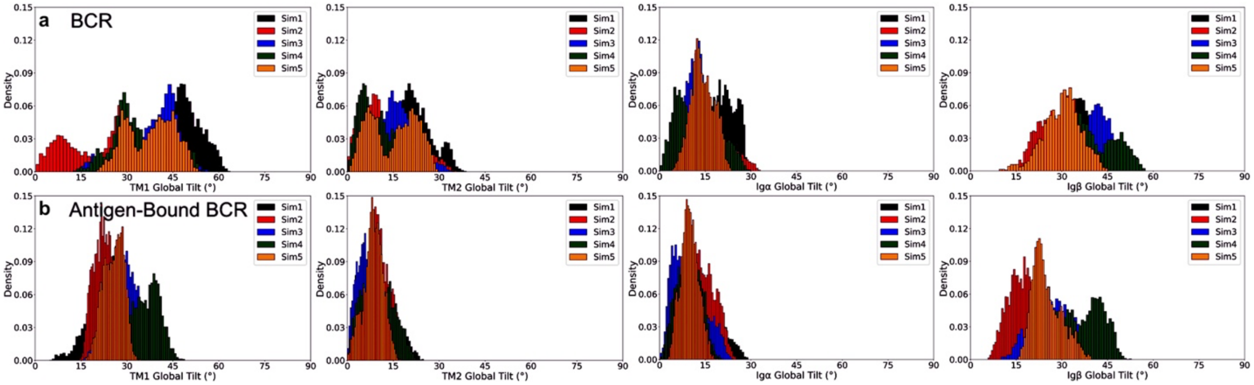

**Supplementary Figure 4. Principal component analysis of the MD simulations of the model CH31 BCR. (a)** Two-dimensional (2D) landscape of the distribution of the BCR conformations along the two primary principal components (PC). A jet color scheme is used to show the distribution density of the BCR conformations. Two low-energy conformational states "S1"-"S2" are observed from the landscape based on their high densities. **(b)** The "S1" state, in which PC1 and PC2 coordinates are 10.3 and 52.6, respectively. **(c)** The "S2" state, in which PC1 and PC2 coordinates are -326.5 and -81.5, respectively. The Fab heavy chains are colored red, Fab light chains are colored blue, Fc domains are colored cyan, transmembrane helices of BCR are colored green, Ig$\alpha$ is colored magenta, and Ig$\beta$ is colored brown. The POPC molecules are colored light blue, POPE are colored light orange, PSM are colored light pink, diacylglycerols (DAGL) are colored pale green, cholesterols (CHOL) are colored wheat, and ceramides (CER3) are colored pale cyan.

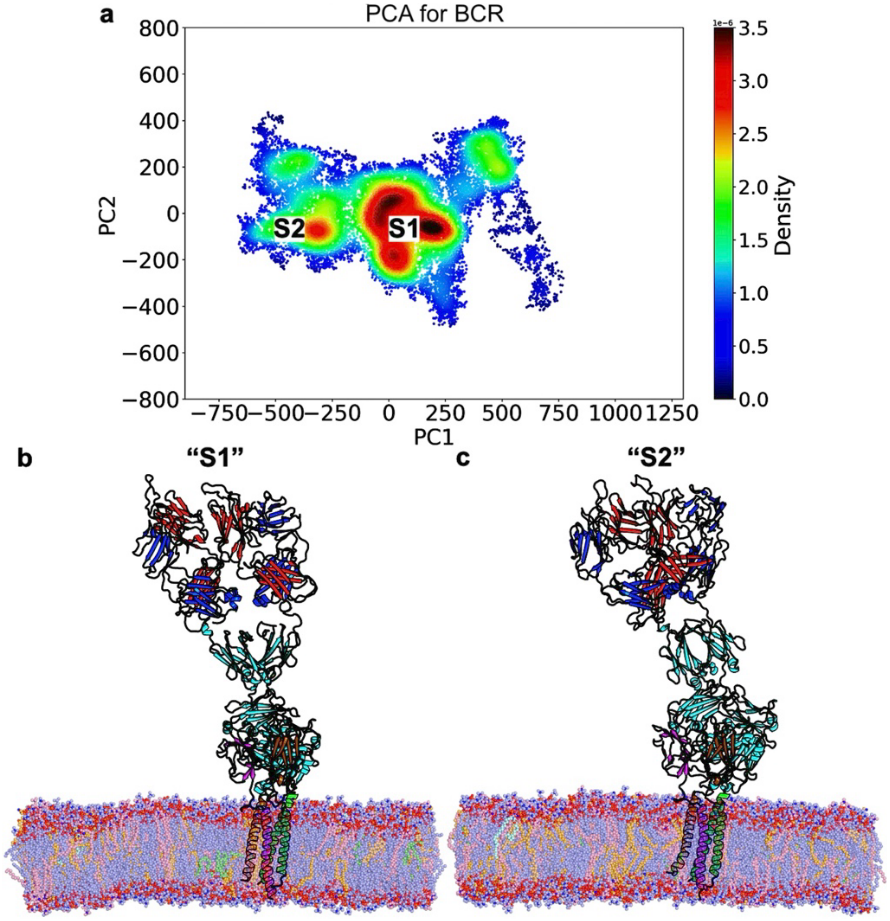

**Supplementary Figure 5. Principal component analysis of the MD simulations of the CH31 BCR bound by HIV-1 Env. (a)** 2D landscape of the distribution of the antigen-bound BCR conformations along the two primary PCs. A jet color scheme is used to show the distribution density of the antigen-bound BCR conformations. Five low-energy conformational states “S1”-“S5” are observed from the landscape based on their high densities. **(d)** The “S1” state, in which PC1 and PC2 coordinates are 13.5 and 83.4, respectively. **(e)** The “S2” state, in which PC1 and PC2 coordinates are 381.4 and -16.1, respectively. **(f)** The “S3” state, in which PC1 and PC2 coordinates are 21.3 and -364.0, respectively. **(g)** The “S4” state, in which PC1 and PC2 coordinates are -400.5 and -36.2, respectively. **(h)** The “S5” state, in which PC1 and PC2 coordinates are 0.0 and 369.1, respectively. The Fab heavy chains are colored red, Fab light chains are colored blue, Fc domains are colored cyan, transmembrane helices of BCR are colored green, Igα is colored magenta, Igβ is colored brown, and the HIV-1 Env is colored orange. The POPC molecules are colored light blue, POPE are colored light orange, PSM are colored light pink, diacylglycerols (DAGL) are colored pale green, cholesterols (CHOL) are colored wheat, and ceramides (CER3) are colored pale cyan.

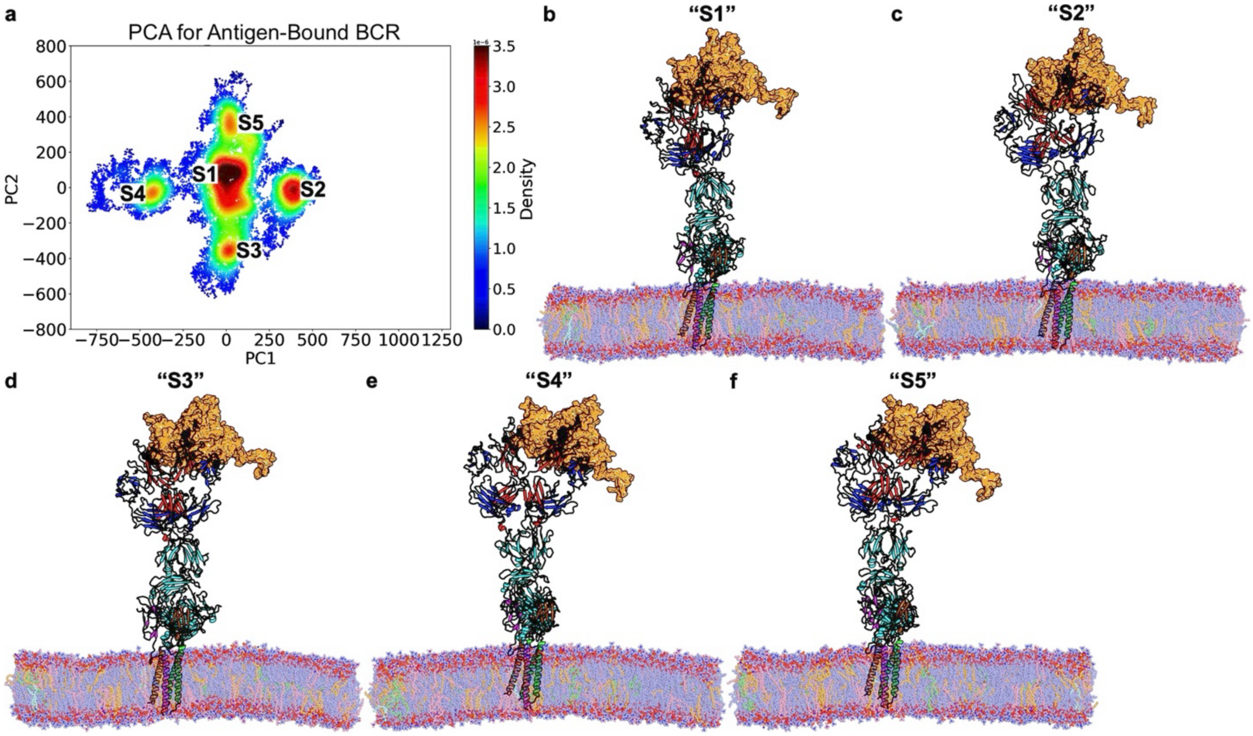

**Supplementary Figure 6. Zoomed in views of the HIV-1 Env bound Fab domain of low-energy conformational states of the antigen-bound BCR. (a)** The "S1" state of the antigen-bound BCR. **(b)** The "S2" state of the antigen-bound BCR. **(c)** The "S3" state of the antigen-bound BCR. **(d)** The "S4" state of the antigen-bound BCR. **(e)** The "S5" state of the antigen-bound BCR. The Fab heavy chains are colored red, Fab light chains are colored blue, Fc domains are colored cyan, and the HIV-1 Env is colored orange.

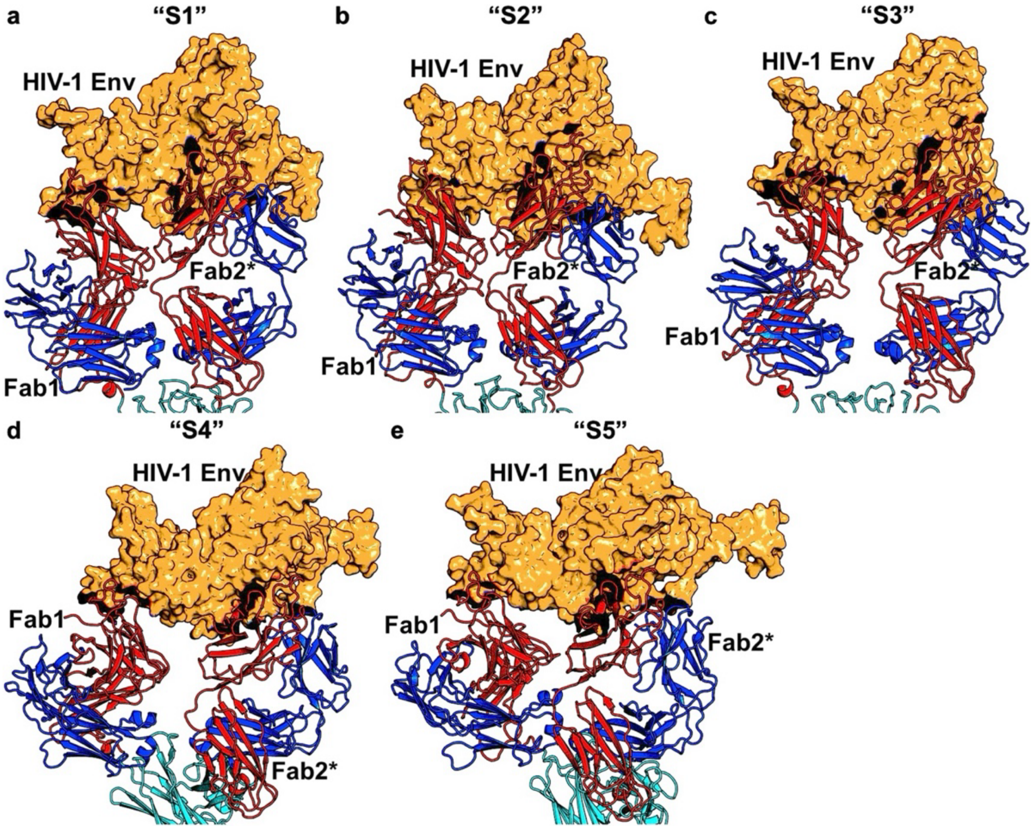

**Supplementary Figure 7. Scheme of the multiscale molecular dynamics (MD) simulation.** All-atom (AA) MD simulations are iterated with long timescale coarse-grained (CG) MD simulations, which are then backmapped to AA simulations to continue the simulations. The CG MD simulations facilitate the lipid mixing in the membranes, while the AA MD simulations refine the protein-protein interactions. The CHARMM36m force field is used for the AA MD simulations, whereas the MARTINI 3 force field is used for the CG MD simulations.

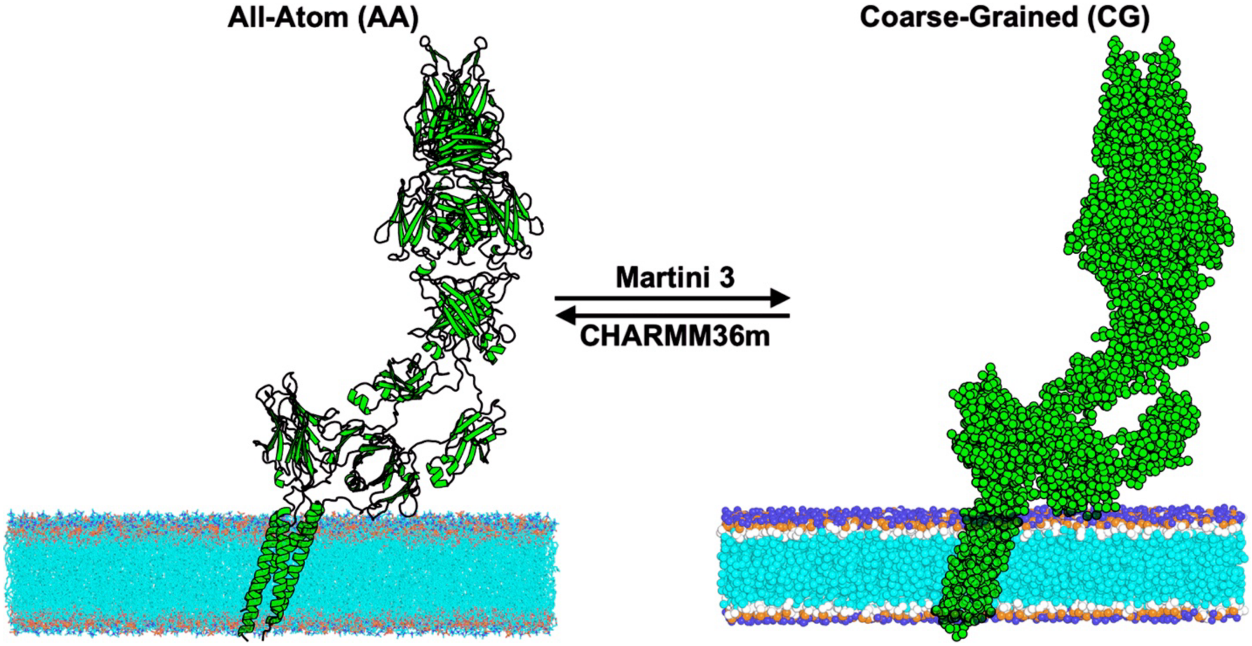

**Supplementary Figure 8. The numbers of surrounding lipid molecules in the initial configurations of the BCR complexes in the complex membrane.** All lipid molecules within 4Å distance of the membrane-peripheral region and membrane-bound helices of the BCR are considered. The bars for the BCR without and with antigen bound are colored blue and orange, respectively.

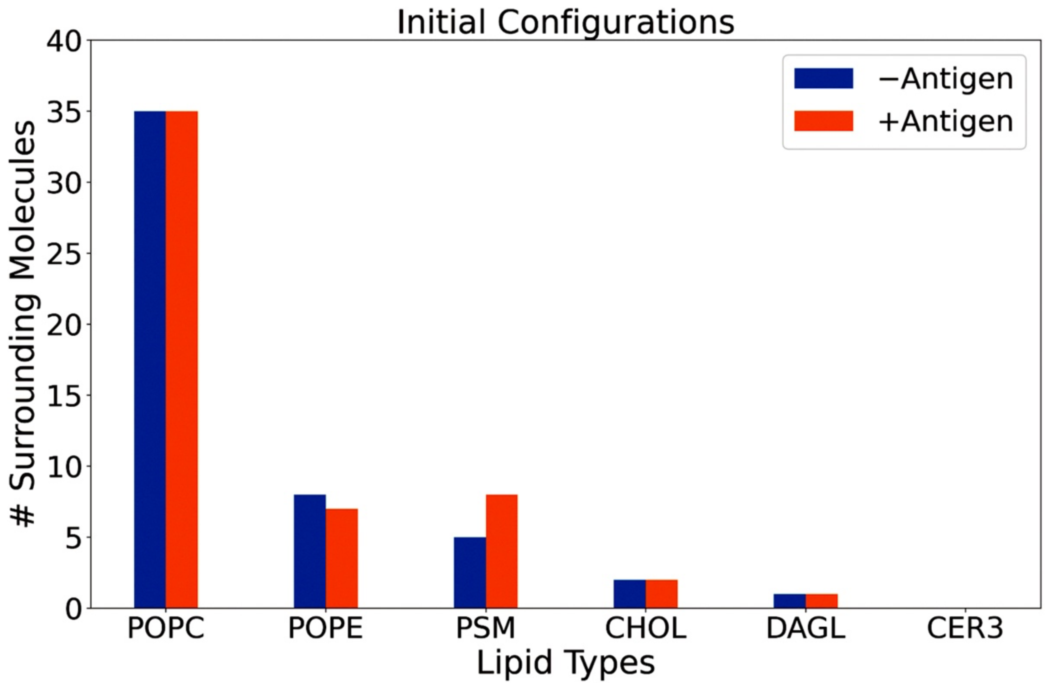

**Supplementary Figure 9.** Diffusions of lipid molecules within the complex membrane during the AA MD simulations of the BCR. A jet color scheme was used to show the speed of lipid diffusion within the membrane.

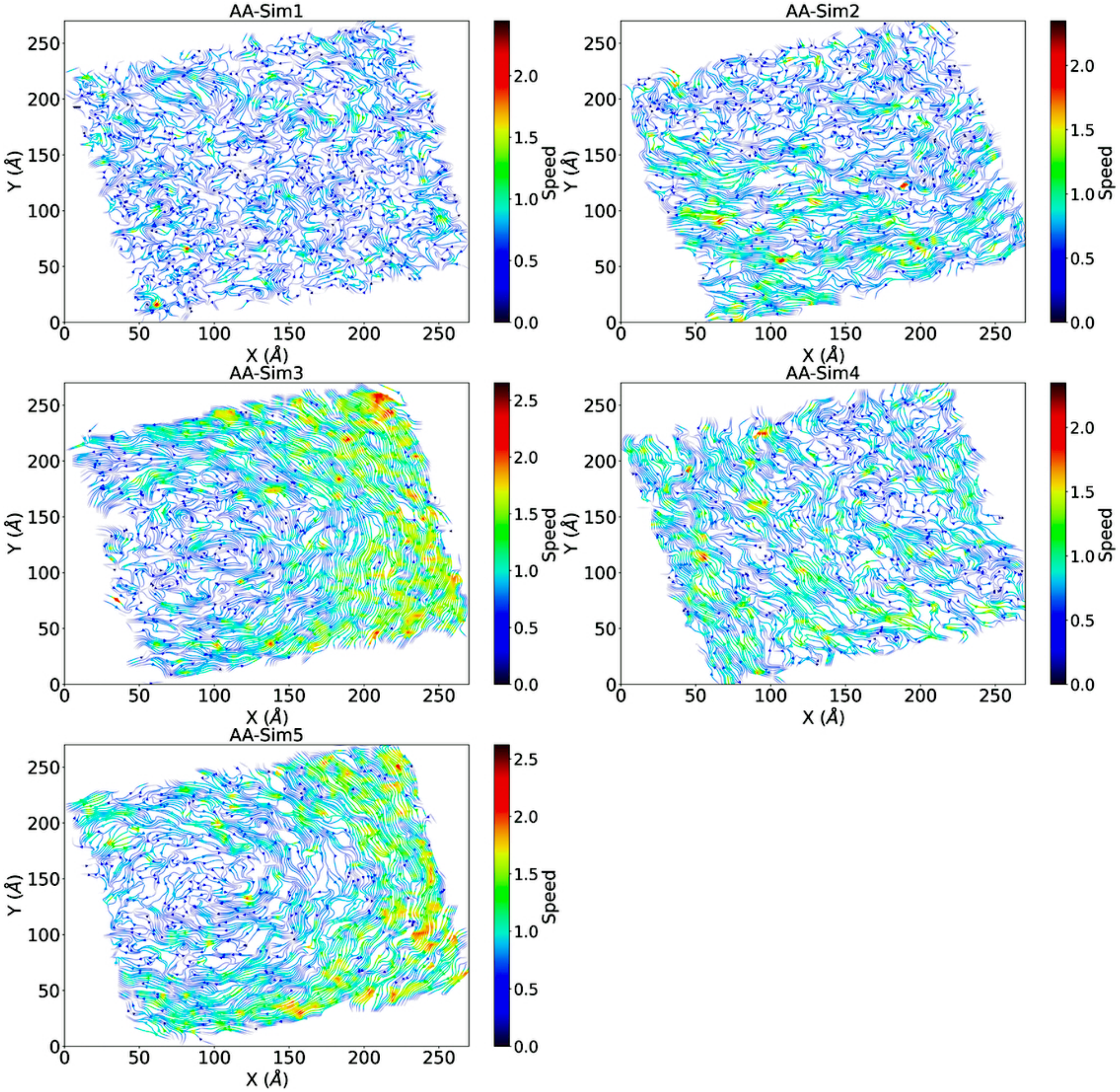

**Supplementary Figure 10.** Diffusions of lipid molecules within the complex membrane during the AA MD simulations of the BCR bound by the HIV-1 Env. A jet color scheme was used to show the speed of lipid diffusion within the membrane.

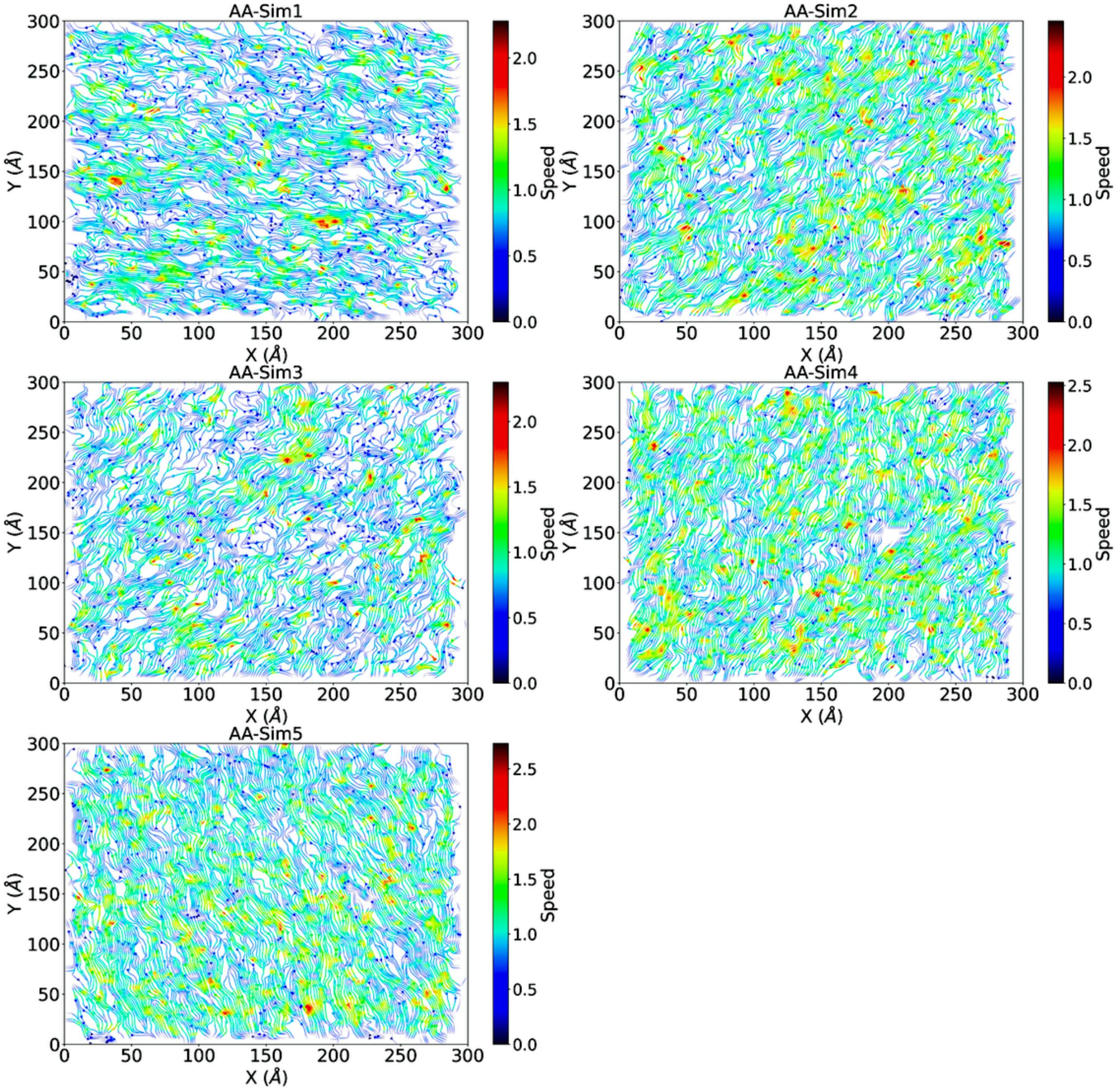

**Supplementary Figure 11.** Diffusions of lipid molecules within the complex membrane during the CG MD simulations of the BCR. A jet color scheme was used to show the speed of lipid diffusion within the membrane.

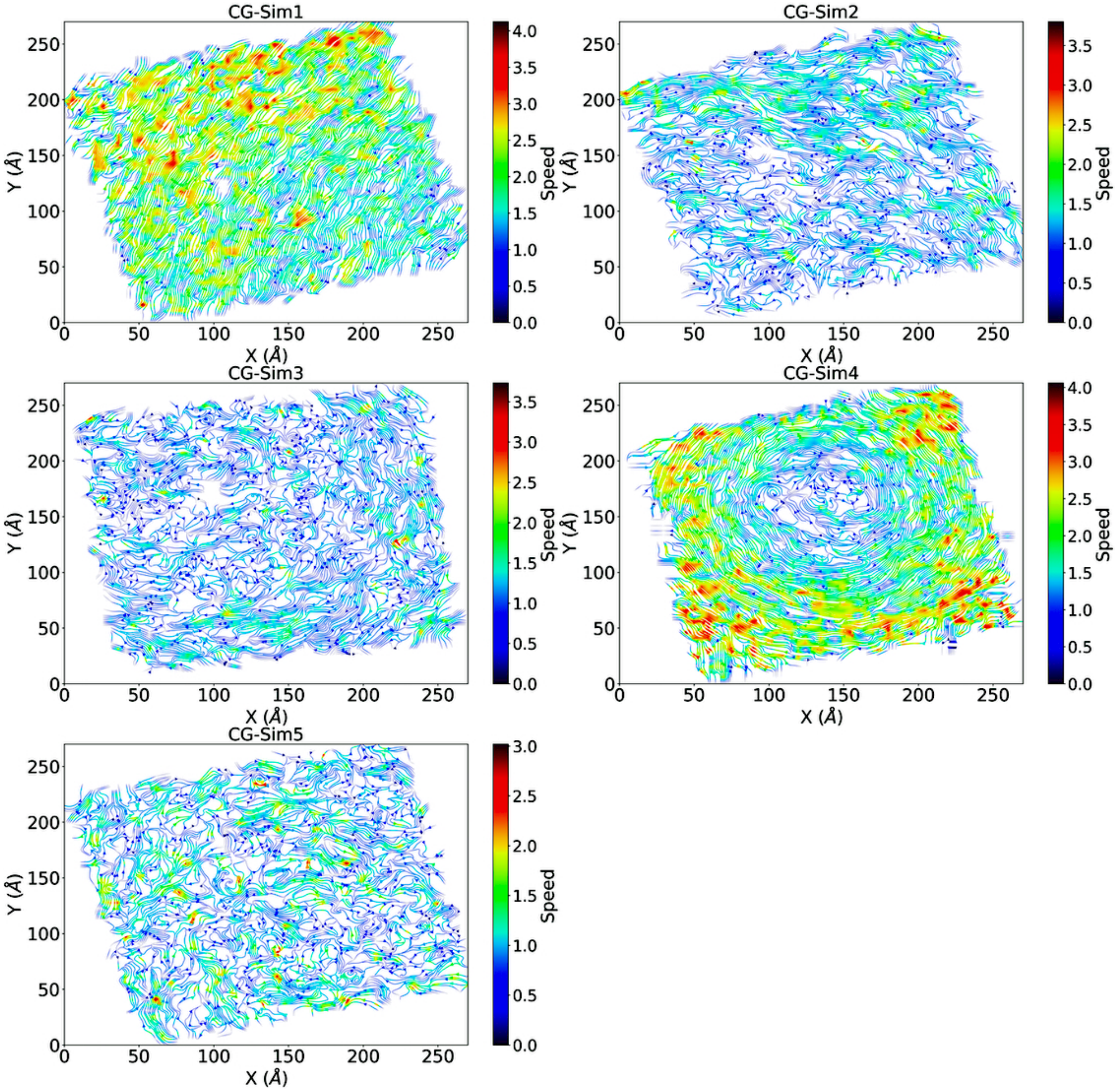

**Supplementary Figure 12.** Diffusions of lipid molecules within the complex membrane during the CG MD simulations of the BCR bound by the HIV-1 Env. A jet color scheme was used to show the speed of lipid diffusion within the membrane.

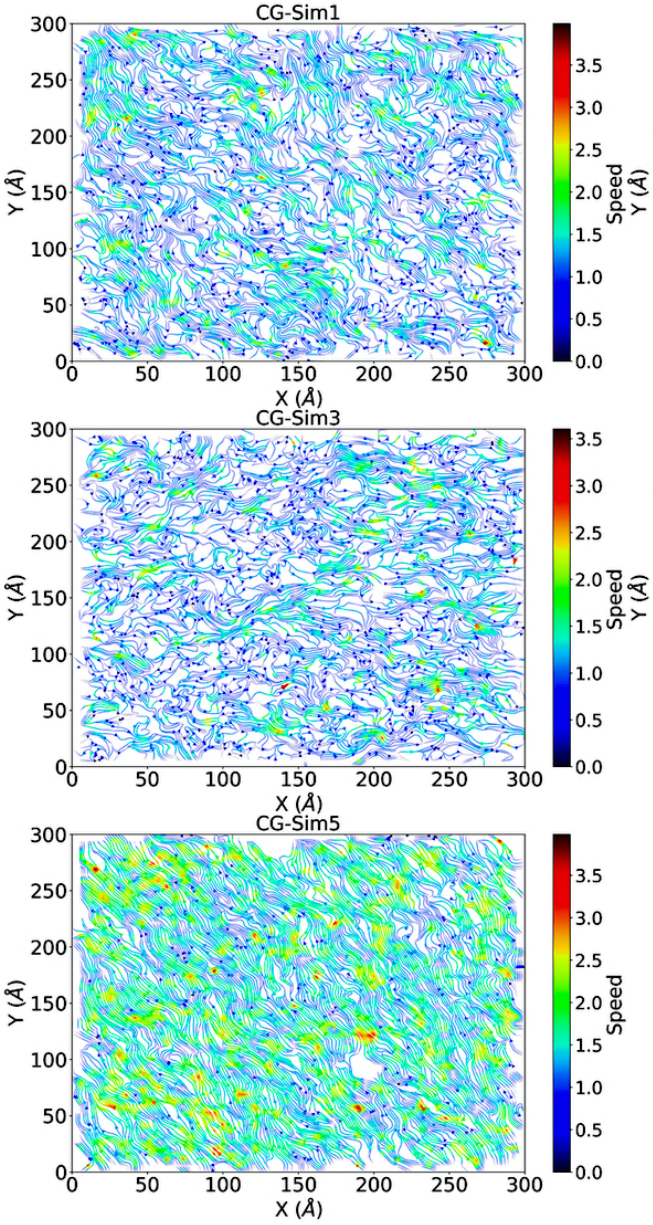

**Supplementary Figure 13. Models of the Igα/Igβ heterodimer and BCR built from their full sequences provided in the 7XQ8 FASTA file by SWISS-MODELLER webserver (a), AlphaFold2-Multimer (b), and AlphaFold3 (c-d).**

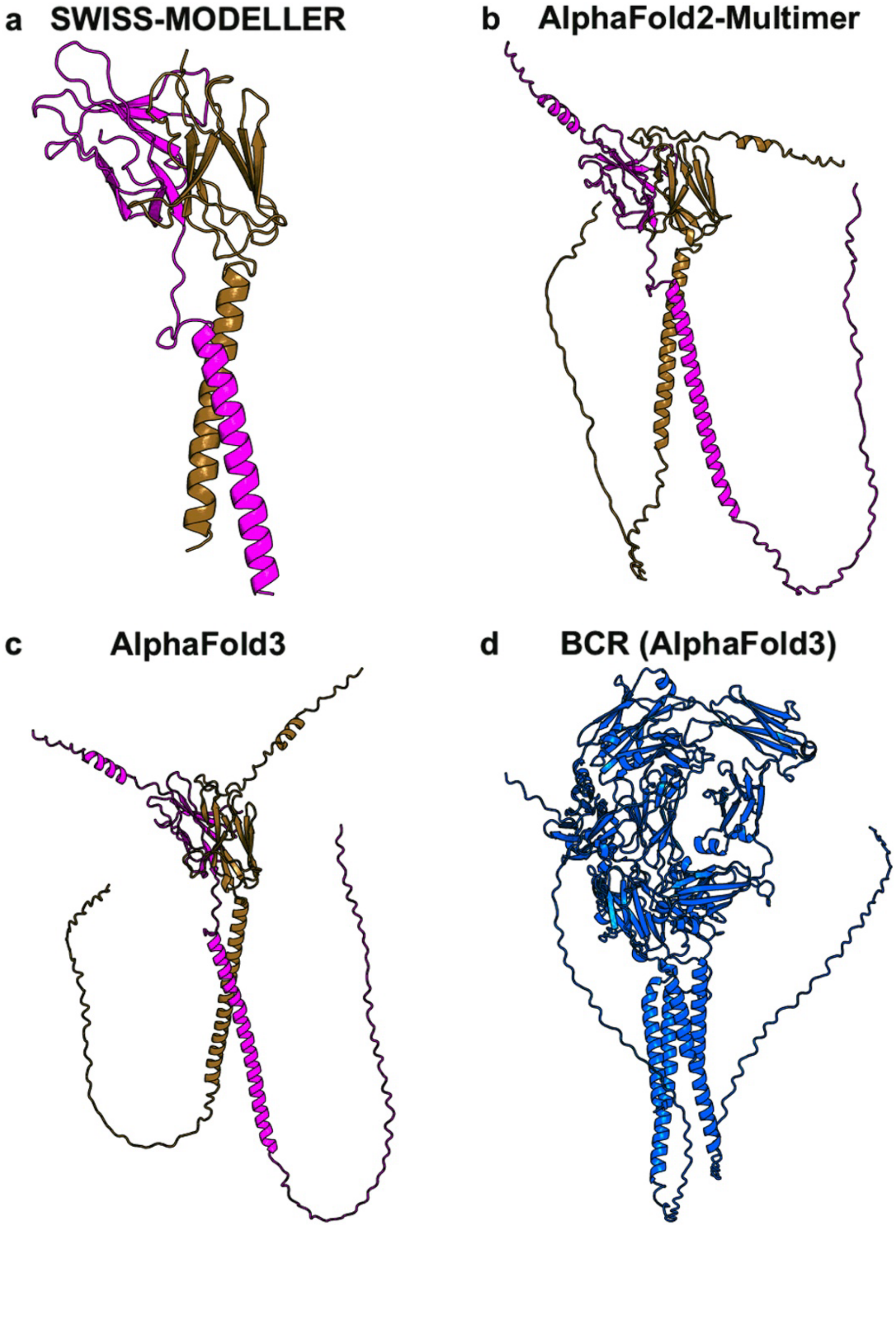
